# Effects of melatonin on germination and amino acid content in different wheat varieties seeds under polyethylene glycol stress

**DOI:** 10.1101/710954

**Authors:** Dongxiao Li, Di Zhang, Hongguang Wang, Haoran Li, Shijia Song, Hongye Li, Ruiqi Li

## Abstract

In this research, the effects of melatonin on germination and amino acid content in wheat (*Triticum aestivum* L.) seeds under polyethylene glycol (PEG-6000) stress were investigated. The results revealed that 10 μM melatonin could aggravate the adverse effects of drought stress on germination percentage, germination index, and germination potential of two wheat varieties (JM22 and HG35). However, 300 μM melatonin could obviously alleviate the negative effect of water stress on germination and increase radicle length, radicle number and plumule length of germinated seeds. Principal component analysis showed that amino acid content apparently changed in germination and the variation amplitude was different due to melatonin concentrations and wheat varieties. Lys content in wheat seed under 300 μM melatonin plus PEG treatment increased compared with that under PEG. Meanwhile, Lys content was significantly and positively correlated with germination percentage, germination index, germination potential, radicle length, and plumule length, respectively. Our results suggested Lys play an important role in the regulation of melatonin on drought resistance of wheat seed during germination. This may be of value for saving water resources in North China Plain.

## Introduction

Drought is a major abiotic stress that lead to great loss in agriculture worldwide. In North China Plain, wheat production has been enduring severely constraint such as yield increasing, quality improving, and groundwater exploitation decreasing. Meanwhile, effective rainfall is scarce for meeting needs of plants during wheat growth stage [1]. As food crop, it is urgent to ensure wheat supply safety for the rapidly increasing world population. Melatonin, a derivative of the essential amino acid tryptophan, can trigger the plant defense responses against adverse environment stress [2–4]. The possible protective roles of melatonin, such as abiotic anti-stressor, biotic anti-stressor, biological rhythm regulator, plant (hormone) regulator and so on, are increasingly curious and researched [5–8].

Previous studies had pointed out that melatonin applied to cucumber seeds could improve their germination rate during chilling stress and water-stress with respect to untreated seeds [4,9]. The pretreatment of seeds with melatonin reduced copper toxicity in red cabbage seedlings *(Brassica oleracea* rubrum) [10]. As is well-known, amino acid metabolism was related and provided nutrient during seed germination [11]. Melatonin share common precursors in the biosynthetic route of tryptophan and tryptamine [12]. Melatonin pretreatment resulted in the osmoprotection through the regulation of proline homeostasis and the enhancement of plant tolerance to drought conditions [13]. A new research reported that melatonin alleviated the inhibitory effects on storage protein degradation of cucumber seed under salt stress, which possible related to amino acid content changing [14].

Additionally, different concentrations melatonin had taken different effects on seeds germination and seedling growth [15]. 1 or 10 μM melatonin could eliminate the inhibitory effect of copper on the fresh weight of seedlings. But 100 μM melatonin had a negative effect on seed germination, seedling grown, and even enhanced the toxic effect of copper [10]. Maize seed priming with 0.8 mM melatonin significantly improved germination energy, germination percentage, proline and total phenolic contents [16]. So far, there are insufficient data on how different concentrations of melatonin affect wheat seed germination by amino acid changing for different wheat varieties.

The main aims of this article were: (1) to investigate the germination characteristics of two wheat varieties at different levels of melatonin over drought conditions; (2) and to evaluate changes in amino acid content during germination of wheat seed. This would help to obtain exact data for further use in wheat yield protection during drought resistance.

## Material and methods

### Tested materials and reagents

The experiment was carried out in a key laboratory of crop growth regulation, Agricultural University of Hebei, Baoding city, China in 2017. Two wheat *(Triticum aestivum* L.) cultivars including the drought-tolerant cultivar ‘Hengguan35’ (HG35) and the irrigated cultivar ‘Jimai22’ (JM22) were used in this study. The HG35 seeds were provided by Dry Land Farming Research Institute of Hebei Academy of Agricultural and Forestry Sciences, whereas the cultivar JM22 seeds were donated by Crop Research Institute, Shandong Academy of Agricultural Sciences. The melatonin and polyethylene glycol 6000 (PEG6000) used in the study were purchased from Bejing Sinopharm Chemical Reagent Co., Ltd.

## Experimental methods

### Preparation of melatonin solution

Firstly, distilled water was used as the basic medium to prepare a 20% PEG solution. Secondly, melatonin was firstly diluted by 1mL 95% ethyl alcohol and then was added to 20% PEG solution to prepare 0, 1, 10, 100 and 300 μmol/L melatonin solutions.

### Experimental treatment

Big and full wheat seeds were selected and sterilized in surface with 70 percent ethanol for two minutes. Then seeds were washed several times with distilled water and were put into the same germination box, in which two-layer filter paper were fully saturated by distilled water (control, CK), polyethylene-glycol solution (20% PEG6000), 20% PEG plus 1 μmol·L^−1^ melatonin solution (1 μM+20% PEG), 20% PEG plus 10 μmol·L^−1^melatonin solution (10 μM+20% PEG), 20% PEG plus 100 μmol·L^−1^ melatonin solution (100 μM+20% PEG), and 20% PEG plus 300μmol·L^−1^ melatonin solution (300 μM+20% PEG), respectively. There were six treatments with three repeats. Each germination box was posed 50 wheat seeds and placed in an incubator at fluctuating day/night temperatures of 20°C/15°C in a light/dark regime. Seed germination was counted daily up to 7 days after placement in the incubator. Water and each concentration solution were complemented into these boxes in time.

### Measurements

The germination percentage was a proportion of emerged-germinating seed in total cultivated seed on 7 days after placement in the incubator.

The germination potential was the germination percentage on 3 days after placement in the incubator.

Root length, sprout length, and radical length of germinated seed were measured 5 repeats with ruler began from 72 hours after placement in the incubator.

Vigor index was measured according to Lu et al. [17].

Vigor index= The germination percentage × radical length

Measurement of amino acid content: Wheat seeds were milled to a fine powder and sifted through 100 mesh. Flour samples (1.00 g) were hydrolyzed for 14 hr at 110°C in the presence of 10 mL of 6 mol·L^−1^ hydrochloric acid. Then it was diluted with water to 10 mL at room temperature. About 1 mL extractions were transferred into centrifuge tube to be concentrated and dried with vacuum chamber. Then the concentrated sample were dissolved with 2 mL hydrochloric acid (0.1 mol·L^−1^). And the dilute sample solution was determinate by reversed-phase high performance liquid chromatograph (HPLC). The HPLC system (Agilent 1200, USA) consisted of a C18 column with 5 μm particle size (4.6 mm inner diameter, 250 mm length, Komati Universal) and a diode array detector. The mobile phase A was prepared by adding acetonitrile. The mobile phase B was prepared by adding acetic acid-sodium acetate buffer (PH 5.25±0.05, glacial acetic acid adjusting) containing 0.03 mol·L^−1^ sodium acetate solution and 0.15% triethylamine. The column temperature was 40°C. The determine wavelength was 360 nm. The flow rate was 1 mL·min^−1^ at ambient temperature. Automatic injection of as many as 42 samples was realized by a G1329A injector with a 10-μL sample loop. The gradient elution procedures were showed at Table 1.

**Table 1.** Gradient elution program.

| Time/min |  | 0 | 10 | 12 | 25 | 30 | 37 | 39 | 45 |
| --- | --- | --- | --- | --- | --- | --- | --- | --- | --- |
| Mobile phase | A | 18 | 18 | 29 | 34 | 55 | 60 | 18 | 18 |
|  | B | 82 | 82 | 71 | 66 | 45 | 40 | 82 | 82 |

Aminoacyl derivatization: 100 μL preparing standard solutions or sample solutions were transferred into 1.5 ml centrifuge tube by adding 200 μL buffered solution (pH 9.0) and 100 μL 2,4-dinitrochlorobenzene (300 mg·mL^−1^, acetonitrile as solvent). The mixture was vortex-mixed for 1 min and extracted in a 90°C thermostatic water bath for 90 min in dark. After the reaction had finished, the resulting solution was added by 50 μL 10%(v/v) acetic acid, adjusted neutral pH value, and diluted the volume to 1mL with distilled water. The solutions were filtered through the organic membrane.

Standard Curve Development: Standard solutions (500 mg·L^−1^) of 17 amino acid including aspartic acid (Asp), glutamic (Glu), histidine (His), serine (Ser), arginine (Arg), glycine (Gly), threonine (Thr), proline (Pro), alanine (Ala), valine (Val), methionine (Met), cysteine (Cys), isoleucine (Ile), leucine (leu), phenylalanine (Phe), lysine (Lys), and tyrosine (Tyr) were prepared by dilution of a stock solution (amino acid standard mixture,10 ml, 0.1mol·L^−1^ HCl). Those were diluted to 5, 50, 125, 100, 150, 200, 300, 400, 500 mg·L^−1^ with 0.1 molL^−1^ HCl. At last, 100μL preparing standard solutions were measured after aminoacyl derivatization performed as above.

### Data processing

All data were run using analysis of variance (ANOVA) with three replicates according to Excel 2010 and SPSS 19.0 (SPSS Inc., Chicago, USA). The Duncan’s new multiple range (DMR) test at 5% probability level was used to test the differences among the mean values.

## Results

### Effects of different concentrations of melatonin on statistical germination

For JM22, germination percentage and germination index decreased significantly by 14.29% and 93.27% under PEG treatments compared with that under control, respectively (Table 2). And 1, 10, 100 μM melatonin plus PEG solution all did not improve germination percentage and germination index. Even 10μM+PEG treatment reduced more on the two values relative to that of PEG. However, germination percentage could be improved significantly under 300μM+PEG treatment comparing with that under other treatments. There was no significant difference on germination percentage between control and 300μM+PEG treatment. But germination index of seed under 300μM+PEG treatment decreased significantly comparing with the control. Additionally, 1 and 10 μL melatonin plus PEG treatments significantly decreased germination potential; whereas it was not changed obviously among other treatments.

**Table 2.**
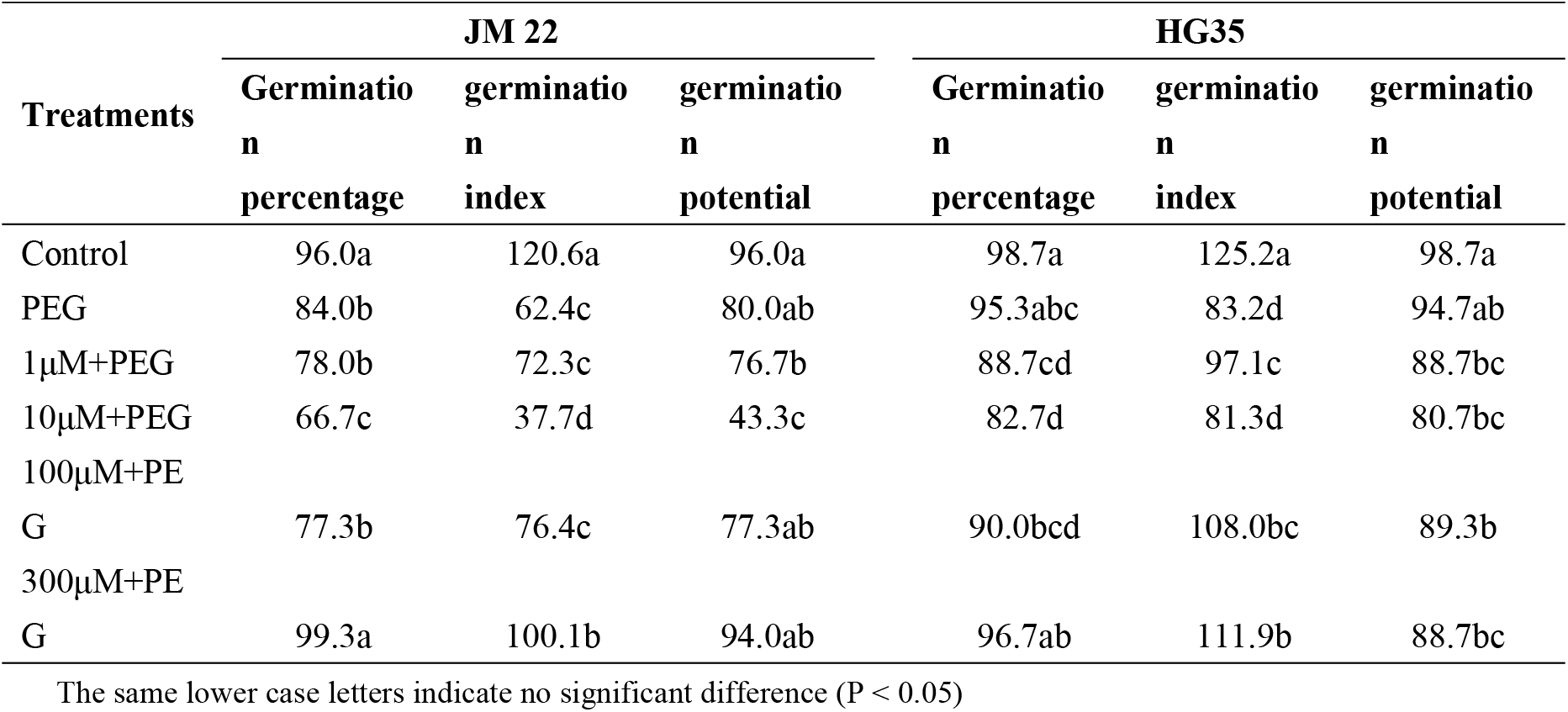
The effect of different treatments on the germination of wheat seeds.

For HG35, compared with the control, germination percentage and germination potential under PEG treatment were both not significantly decreased; but germination index decreased by 50.48% (P<0.05). 10μM+PEG treatment further decreased germination percentage significantly. Compared with PEG, germination index under 100μM+PEG and 300μM+PEG treatments increased by 29.81% and 34.50%, respectively (P<0.05).

### Effects of different concentrations of melatonin on morphological characters of germination

In this study, radicle length and radicle number of two wheat cultivars were both increased gradually from 72 hours to 144 hours after germination (Fig 1). For JM22, radicle length under PEG and 10μM+PEG treatments decreased by 131.00% and 189.00% on average compared with the control. No obvious change had been found on that under 1μM+PEG and 100μM+PEG treatments. However, 300μM+PEG treatment resulted in an increase of 79.01% on radicle length compared with that under PEG treatment. Compared with the control, radicle number under other treatments were decreased significantly; and the least value under 10μM+PEG treatments decreased by 79.6% (P<0.05). Under 300μM+PEG treatment, radicle number increased by 24.41% compared with that under PEG treatment. And the plumule length was decreased significantly under all other treatments compared with the control. There was hardly obvious improving effect of melatonin on plumule length under PEG treatment.

**Fig 1.**
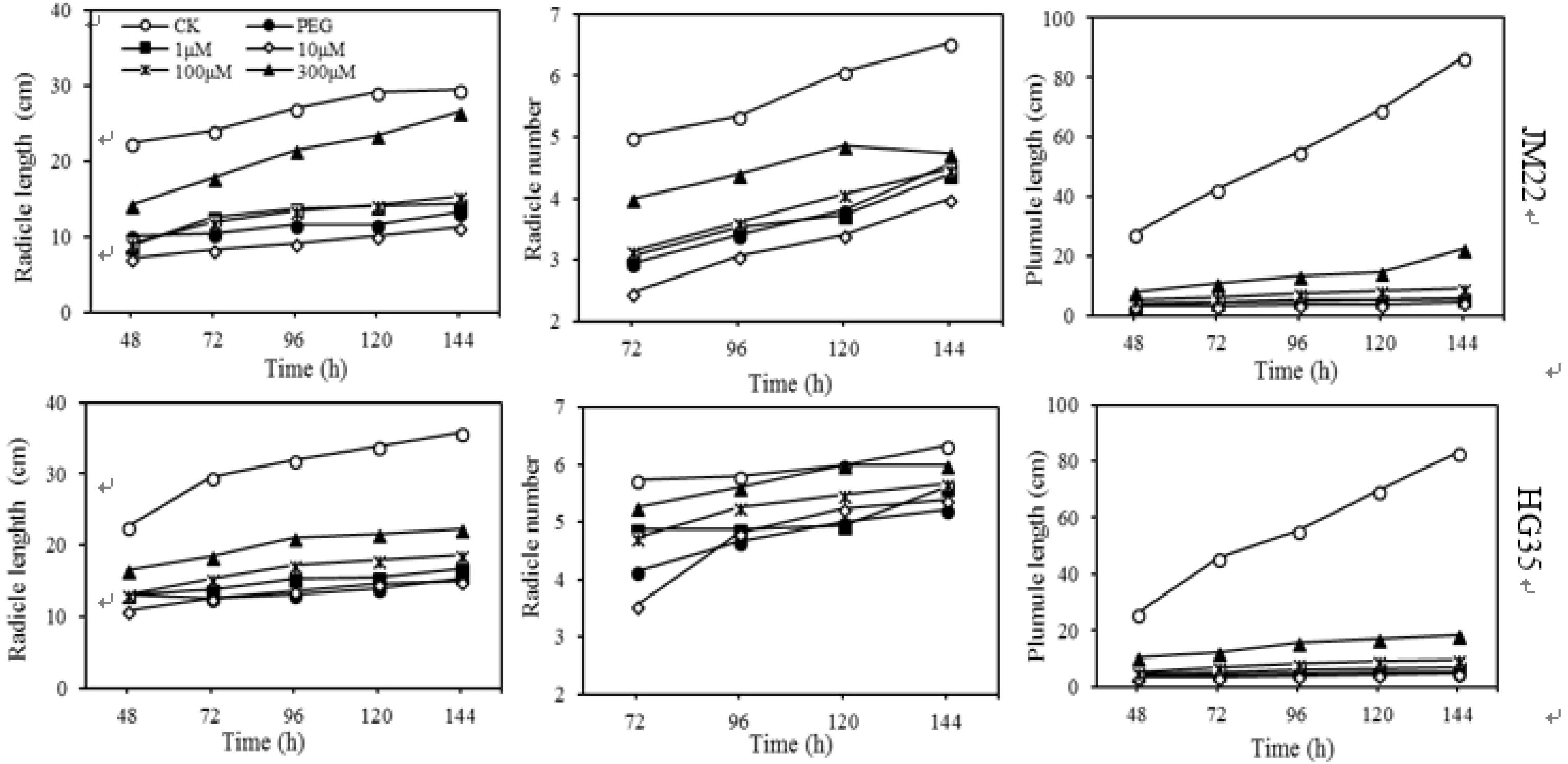
Effects of different concentrations of melatonin on radicle length, radicle number, and plumule length of JM22 and HG35. Values are means of three biological replications.

For HG35, compared with the control, radicle length and plumule length were both decreased significantly under all other treatments. Radicle length of wheat seed under 300 μM melatonin plus PEG treatment decreased by 85.72% compared with the control; but it increased by 20.65% compared with PEG treatment. There was no significant difference on radicle number between control and 300μM+PEG treatment. The changing trend of plumule length was similar to that of JM22. Additionally, as a drought-resistant variety, radicle length and radicle number of HG35 were both higher than that of JM22 under PEG treatment.

### Effects of different concentrations of melatonin on amino acid content

The results showed the variations among all different treatments (Fig 2). Principal component analysis extracted two major components that accounted cooperatively for 74.8% of the variance in the data set. Principal component 1 (PC1, X-axis) explained 43.8% of the variation among the individual samples, principal component 2 (PC2, Y-axis) explained 31.0% of the variation. When seed germinated, the amino acid content of both cultivars shift greatly along PC2 axis. This meaning that amino acid content apparently changed in germination. When seed germinated with PEG-6000, the amino acid content of both cultivars goes down along PC2 axis; moreover, HG35 and JM22 were separated with each other. This indicated a genotypic variation of amino acid content in seed germination with PEG-6000. This may be the reason why seed germination of HG35 was better than that of JM22 under water stress condition.

**Fig 2.**
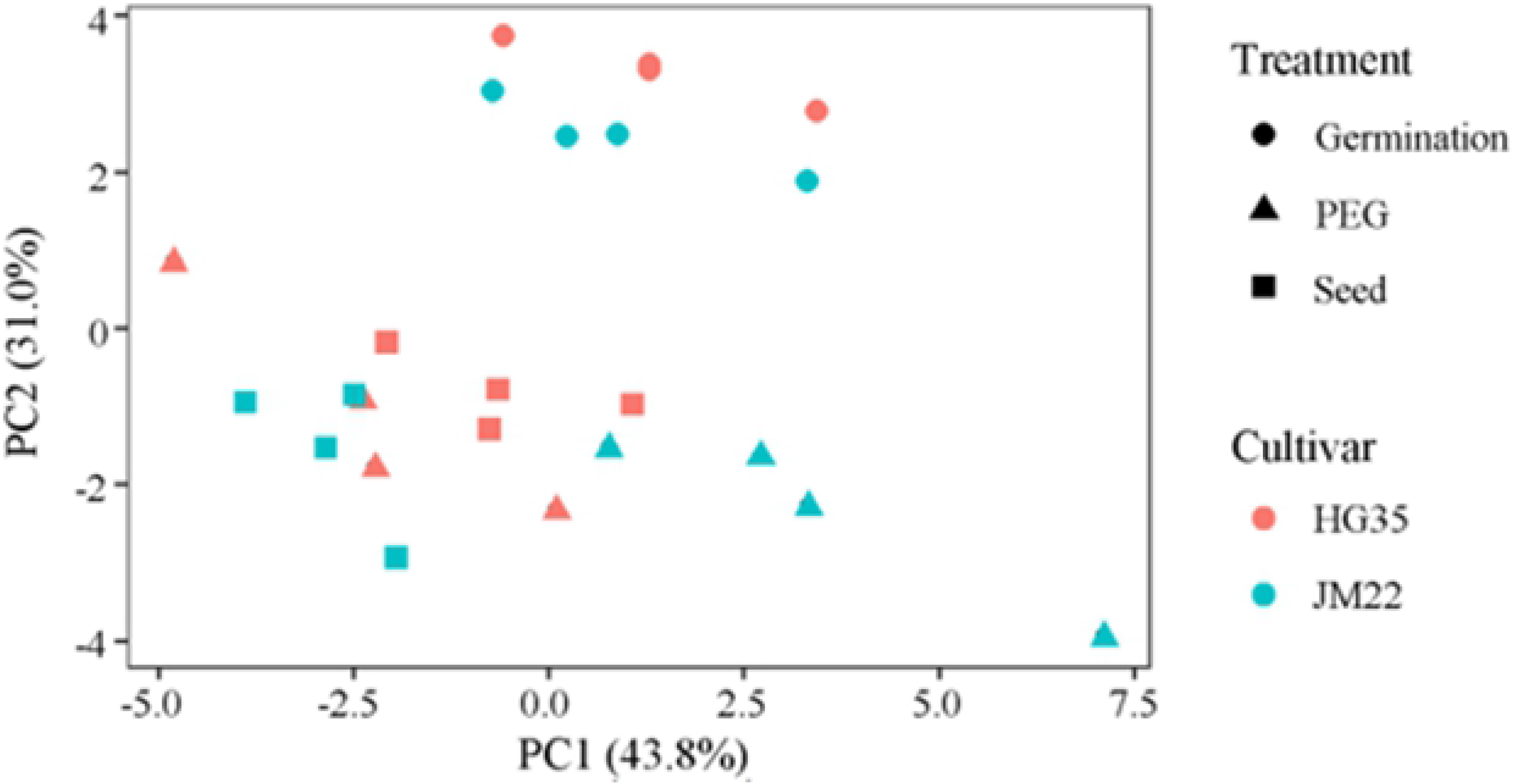
Principal component analysis of amino acid for different wheat cultivars. Germination represent dry seeds germinated by distilled water; PEG represent dry seeds germinated by 20% PEG; Seed represent dry seeds.

Glu content was the highest in wheat seed of both cultivars, followed by Pro content. After seed germinated, Glu and Pro content reduced by 17.16 mg kg^−1^ and 4.41 mg kg^−1^ for JM22; and those reduced by 12.96 mg kg^−1^ and 3.56 mg kg^−1^ for HG35, respectively (Table S1). This indicated that Glu and Pro could be the source of nitrogen during quiescent dry seed resuming metabolic activity. After germinated, Lys and Tyr contents in CK seeds both increased by 129.22% and 130.77% for JM22; and those increased by 71.85% and 98.59% for HG35, respectively. But comparing with CK, Lys and Tyr contents in seeds under other treatments were all decreased at different levels; Glu and Pro contents were reversely increased (Fig 3). The variation amplitude of amino acid content was different due to melatonin concentrations and wheat varieties. For JM22, comparing with CK, Lys and Tyr content reduced by 48.44% and 36.67% in seed under PEG; melatonin plus PEG treatments raised the reduction except 300 μM melatonin reducing Lys content by 37.96%. Compared with CK, Glu and Pro content increased by 44.41% and 29.52% in seed under PEG; and the two indexes increased by 74.40% and 48.19% on average under PEG plus low concentration melatonin (1 and 10 μM). For HG35, comparing with CK, Lys and Tyr content reduced by 62.72% and 53.19% under PEG; Lys content reduced by 57.97% and 39.87% under melatonin plus 100 μM and 300 μM treatments, respectively. On the same situation, Glu and Pro content increased by 51.97% and 55.27% in seed under PEG; the increasing amplitude of the two indexes decreased under PEG plus different concentrations melatonin treatments.

**Fig 3.**
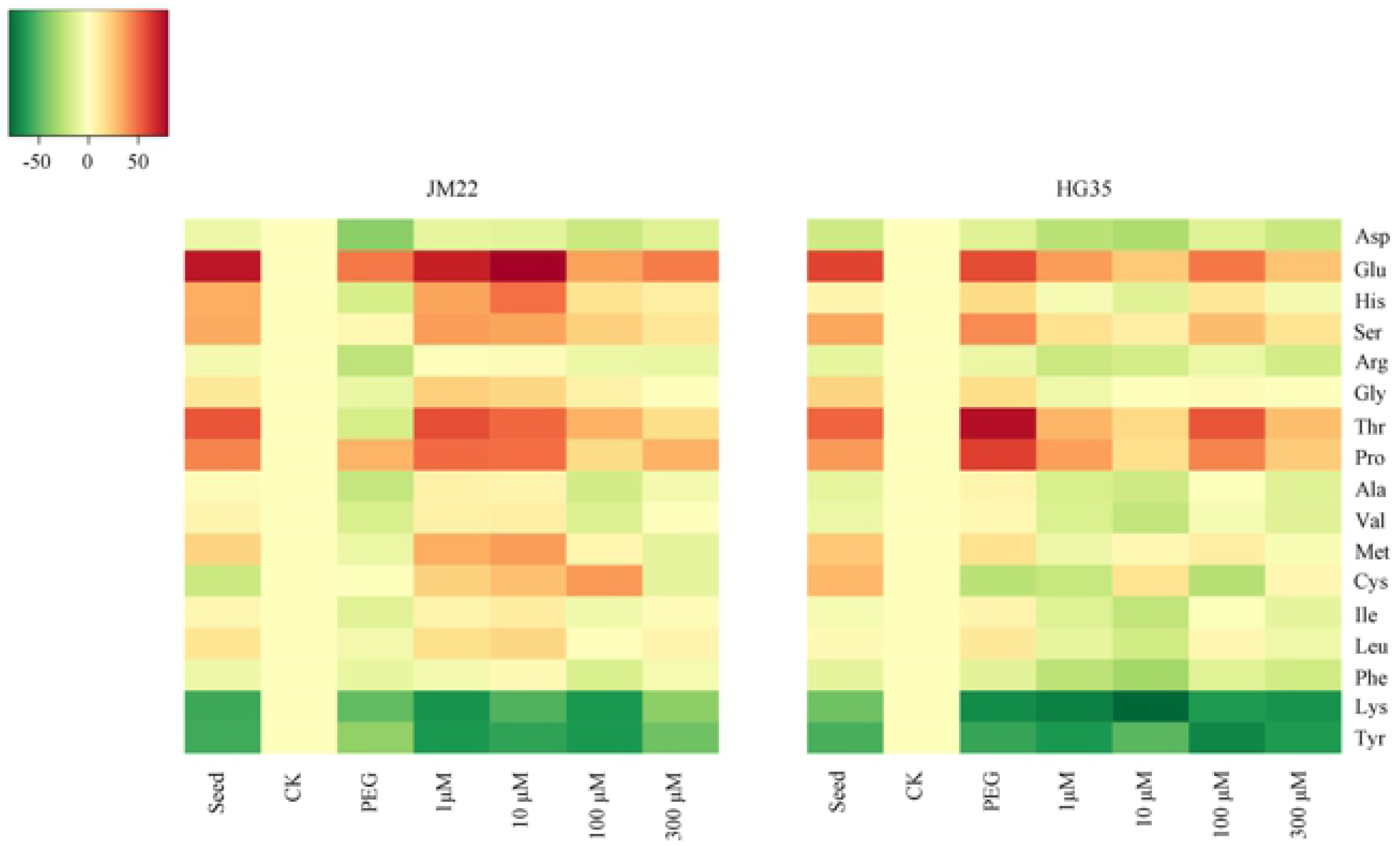
The cluster heatmap of amino acid content differences of two wheat varieties during germination under different treatments. The graph’s horizontal axis shows different treatments (seed represent no-germinated seeds; CK represent seeds germinated by distilled water; PEG represent seeds germinated by 20%PEG; 1μM, 10μM, 100μM, and 300μM, represent seeds germinated by 1μM, 10μM, 100μM, and 300μM melatonin plus 20%PEG, respectively), and the vertical axis shows different amino acid. Color gradients represent the differences value of amino acid contents under other treatments compared with that of CK.

Compared with PEG treatment, the contents of Glu, Met, Cys, and Tyr in JM22 seed decreased under 300μM melatonin plus PEG treatment, but other amino acid contents increased (Fig 3 and Table 1S). Cys and Lys content in HG35 seed under 300μM melatonin plus PEG treatment was higher than that of PEG treatment, whereas other components was lower.

### Correlation between amino acid content and morphological indexes

Correlation coefficients revealed that Asp content was significantly and positively correlated with germination index, radical length, and radical number (Table 3). Glu content was negatively correlated with all morphological indexes of germination, especially plumule length. After seed germinated, Glu and Pro content reduced significantly (Table S1 and Fig 3). This indicated that Glu and Pro possibly provided amino (NH_4_^+^) to other amino acid by aminotransferase when seed germinated. Phe content was significantly and positively correlated with radical number. And Lys content was significantly and positively correlated with germination percentage, germination index, germination potential, radicle length, and plumule length, respectively. This suggested that Lys played an important role in wheat germination and had an obvious interaction with melatonin.

**Table 3.**
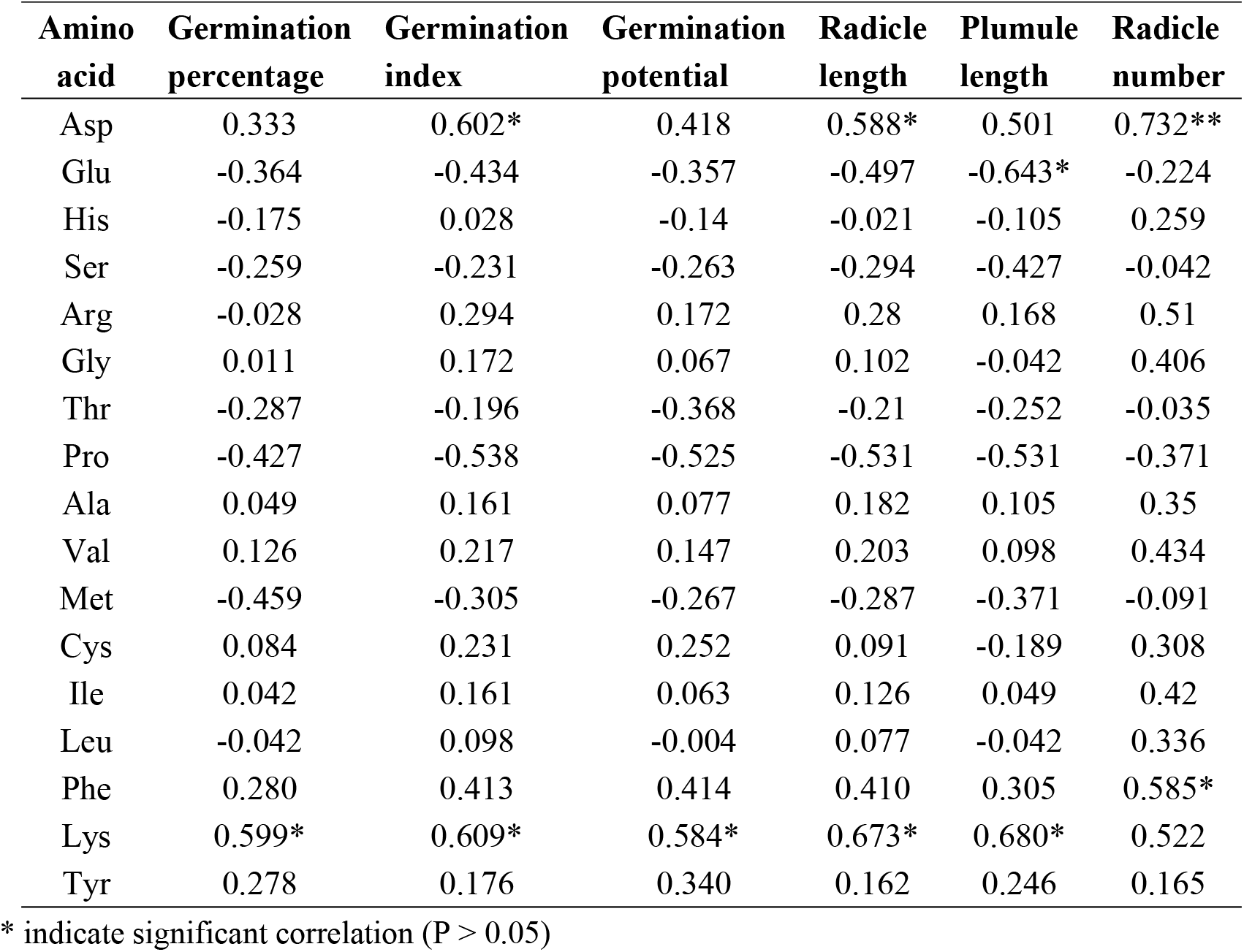
The correlation analysis of variance between morphological index of germination and 17 amino acids.

| Amino acid | Germination percentage | Germination index | Germination potential | Radicle length | Plumule length | Radicle number |
| --- | --- | --- | --- | --- | --- | --- |
| Asp | 0.333 | 0.602* | 0.418 | 0.588* | 0.501 | 0.732** |
| Glu | -0.364 | -0.434 | -0.357 | -0.497 | -0.643* | -0.224 |
| His | -0.175 | 0.028 | -0.14 | -0.021 | -0.105 | 0.259 |
| Ser | -0.259 | -0.231 | -0.263 | -0.294 | -0.427 | -0.042 |
| Arg | -0.028 | 0.294 | 0.172 | 0.28 | 0.168 | 0.51 |
| Gly | 0.011 | 0.172 | 0.067 | 0.102 | -0.042 | 0.406 |
| Thr | -0.287 | -0.196 | -0.368 | -0.21 | -0.252 | -0.035 |
| Pro | -0.427 | -0.538 | -0.525 | -0.531 | -0.531 | -0.371 |
| Ala | 0.049 | 0.161 | 0.077 | 0.182 | 0.105 | 0.35 |
| Val | 0.126 | 0.217 | 0.147 | 0.203 | 0.098 | 0.434 |
| Met | -0.459 | -0.305 | -0.267 | -0.287 | -0.371 | -0.091 |
| Cys | 0.084 | 0.231 | 0.252 | 0.091 | -0.189 | 0.308 |
| Ile | 0.042 | 0.161 | 0.063 | 0.126 | 0.049 | 0.42 |
| Leu | -0.042 | 0.098 | -0.004 | 0.077 | -0.042 | 0.336 |
| Phe | 0.280 | 0.413 | 0.414 | 0.410 | 0.305 | 0.585* |
| Lys | 0.599* | 0.609* | 0.584* | 0.673* | 0.680* | 0.522 |
| Tyr | 0.278 | 0.176 | 0.340 | 0.162 | 0.246 | 0.165 |

\* indicate significant correlation ( $P > 0.05$ )

## Discussion

Different concentrations of melatonin have taken different effects on crop growth [18–19]. Melatonin (0-1 μM) promoted the root growth; but the higher melatonin concentrations (5-10 mM) inhibited root growth and chlorophyll concentration of cherry rootstock PHL-C [20]. In the present study, 10 μM melatonin could aggravate drought stress effect on wheat seed germination under PEG condition. It has been reported that exogenous melatonin with low concentration (1 μM and 10 μM) inhibited leaf physiology by communication of abscisic acid and hydrogen peroxide [15]. And this study also showed that 300 μM melatonin could obviously alleviate the drought stress effect on seed germination of wheat. This was in accordance with that of Cui and others [21], reporting that melatonin significantly affected the expression of glycolytic proteins, modulated electron transport in the respiratory chain, and improved energy production.

Seed germination is an important biological and dynamic process with mobilization of the major storage reserves. Under water stress, proline content in wheat seed significantly increased [22]. Proline as organic substances can regulate the plasma osmotic potential, and protect the enzymes and plasma membranes under water stress condition [23]. A low concentration of melatonin promoted the synthesis of glycine and succinyl-CoA to influence porphyrins synthesis; high concentration of melatonin lead to proline and carbohydrate synthesis for osmoregulation [20]. Melatonin could also accelerate the metabolic flow from the precursor amino acids arginine and methionine to polyamines, which mitigate salt stress on wheat seedling [24]. These suggested that melatonin had directly and indirectly influenced on amino acid content. In this study, Pro and Glu content increased in wheat seed under PEG stress. Low concentration of melatonin (1 and 10 μM) improved the increasing range of Pro, Glu and Gly content in seed of JM22 under water stress, which was partially conformed to Sarropoulou et al. [20]. However, melatonin decreased the increasing range of Pro and Glu content in seed of HG35. Results showed different drought-resisting ability of two wheat varieties.

During germination, amino acids stored in wheat seeds as storage protein are decomposed by hydrolysis [25]. Under abiotic stress, such as salt and water deficiency, free amino acids would play regulation roles as the most hydrophilic and somatically active compounds [26]. Hartmann et al. [27] reported a pathogen-inducible Lys catabolic pathway in plant that generated N-hydroxypipecolic acid as a critical regulator of systemic acquired resistance to pathogen infection. In this study, Asp, Glu, Phe, and Lys contents were all correlated significantly to one or more indexes of wheat germination. Remarkably, Lys content was significantly and positively correlated with germination percentage, germination index, germination potential, radicle length, and plumule length. A new report showed that histone deacetylase 14, playing a significant role in deacetylation of lysine on histone, was involved in melatonin biosynthesis of *Arabidopsis* thaliana [28]. And future studies should be focused on the interaction between melatonin and prominent amino acid such as lysine changing in different wheat varieties seeds under water stress.

## Conclusion

In summary, our data revealed that exogenous melatonin with different concentrations had different effects on germination morphological indexes of winter wheat. 300 μM melatonin could obviously alleviate the adverse effect of drought stress on germination of wheat seed. This was accompanied by an increase in lysine content, which was significantly and positively correlated to seed gemination indexes. This suggested Lys may play an important role in regulating drought resistance of wheat seed treated by melatonin. Also, there was genotypic difference in drought resistance due to difference in amino acid content and changing amplitude during wheat germination.

## Supporting Information

S1 Table. The objective value corresponding to Fig 3.

## Acknowledgements

This work was financially supported by the National System of Modern Agriculture Industrial Technology Project (CARS-03-05), the National Key Research and Development Program of China (2017YFD0300909), the Scientific Research Project of Hebei Education Department (QN2019046), and the Hebei Province Natural Science Foundation for Youth (C2019204358).

## Author contribution statement

**Data curation:** Di Zhang, Hongguang Wang, Haoran Li

**Methodology:** Shijia Song, Hongye Li

**Supervision:** Ruiqi Li

**Writing – original draft:** Dongxiao Li, Di Zhang

**Writing – review & editing:** Dongxiao Li

---

**Table S1 Effect of different concentrations of melatonin on amino acid content in winter wheat under PEG treatments**

